# Schwann cell p75NTR sustains persistent pain downstream to NGF through ROS-dependent TRPA1 signaling

**DOI:** 10.64898/2026.08.10.743924

**Authors:** Matilde Marini, Alice Papini, Martina Chieca, Elisa Bellantoni, Giovanna Pivotto, Lucrezia Timotei, Gaetano De Siena, Maryam Raeispour, Alexandra Dimitrova, Lorenzo Bonacchi, Gabriele Ferroni, Irene Scuffi, Natália Gabriele Hösch, Sabrina Qader Kudsi, Francesco De Logu, Romina Nassini

## Abstract

Nerve growth factor (NGF) is a key mediator of pain through activation of the high-affinity tropomyosin receptor kinase A (TrkA) and the low-affinity neurotrophin receptor (p75NTR). Although neuronal TrkA signaling is well established, the contribution of non-neuronal cells to NGF-dependent pain remains unclear. Here, we show that NGF and its precursor proNGF engage distinct cellular mechanisms. Intraplantar NGF induced acute nociception, heat hyperalgesia, mechanical allodynia, and cold hypersensitivity, whereas cleavage-resistant proNGF selectively evoked mechanical allodynia and cold hypersensitivity. Pharmacological and cell-specific genetic approaches demonstrated that acute nociception and heat hyperalgesia require neuronal TrkA, whereas mechanical and cold hypersensitivity depend on p75NTR activation in Schwann cells. In Schwann cells, NGF and proNGF induced p75NTR-dependent calcium release, followed by TRPA1 activation, mitochondrial ROS production, and NOX1-dependent oxidative amplification. Inhibition of ROS or TRPA1, or Schwann cell-specific *Trpa1* deletion, markedly reduced mechanical allodynia and cold hypersensitivity without affecting acute nociception or heat hyperalgesia. These findings identify a Schwann cell p75NTR-ROS-TRPA1 pathway sustaining persistent pain and highlight non-neuronal p75NTR signaling as a potential therapeutic target.

## INTRODUCTION

Nerve growth factor (NGF) was first described in 1951 as a critical regulator of sensory and sympathetic neuronal differentiation during development (Sofroniew *et al*, 2001). It exerts essential functions in both the central (CNS) and peripheral nervous systems (PNS). In the CNS, NGF regulates the number of choline acetyltransferase-positive neurons and contributes to learning and memory processes (Huh *et al*, 2008; Niewiadomska *et al*, 2011). In PNS, it controls the size and cellular composition of dorsal root ganglia (Yip *et al*, 1984). The development of peptidergic small- and medium-diameter DRG neurons is impaired in NGF-null mice and in mice lacking TrkA receptor (Crowley *et al*, 1994; Smeyne *et al*, 1994).

NGF signals through two receptors: the high-affinity tropomyosin receptor kinase A (TrkA) and the low-affinity p75 neurotrophin receptor (p75NTR), the latter also capable of binding the precursor form, proNGF (Minnone *et al*, 2017). Beyond their trophic functions, NGF and its receptors NGF are key mediators in pain signaling (Barker *et al*, 2020). NGF contributes to inflammatory pain by sensitizing primary sensory neurons, promoting sympathetic nerve sprouting, and accumulating at sites of tissue injury, thereby contributing to persistent pain hypersensitivity (Longo *et al*, 2013; Woolf *et al*, 1994). The NGF-TrkA interaction forms a receptor-ligand complex that undergoes endocytosis and retrograde transport, modulating nuclear transcription (Grimes *et al*, 1997; Grimes *et al*, 1996; Peach *et al*, 2024), which results in the upregulation of pro-nociceptive mediators including substance P and CGRP released by nociceptors following noxious stimulation (Donnerer *et al*, 1992; Lewin & Mendell, 1993). In parallel, activation of p75NTR can sensitize primary sensory neurons by increasing ceramide production (Dobrowsky *et al*, 1994), enhancing ASIC3 expression (Mamet *et al*, 2002), upregulating bradykinin binding sites (Rueff *et al*, 1996), and promoting substance P synthesis (Lindsay & Harmar, 1989). Despite intense preclinical and clinical efforts targeting NGF signaling, therapeutic strategies aimed at modulating this pathway have not yet achieved sustained clinical approval, highlighting the need for a deeper mechanistic understanding of NGF-dependent pain pathways.

Chronic pain affects more than 30% of people worldwide. Although current treatments, including non-steroidal anti-inflammatory drugs (NSAIDs), opioids, and antidepressants, may provide symptomatic relief, their long-term use is often limited by significant adverse effects (Cohen *et al*, 2021). While nociceptors are traditionally considered the primary drivers of pain transmission to CNS, accumulating evidence indicates that non-neuronal cells critically contribute to the initiation and persistence of chronic pain through bidirectional interactions with sensory neurons (Inoue & Tsuda, 2018; Ji *et al*, 2013). We recently demonstrated that Schwann cells, classically known for their trophic and myelinating functions, actively participate in migraine (De Logu *et al*, 2022), inflammatory pain(Nassini *et al*, 2025) cancer pain (Landini *et al*, 2023), and endometriosis-associated pain (Titiz *et al*, 2024). Notably, p75NTR is highly expressed in Schwann cells, where it regulates myelination, survival, and migration during nerve regeneration (Goncalves *et al*, 2020).

Here we observed that intraplantar administration of NGF and proNGF induced mechanical allodynia and cold hypersensitivity, whereas only NGF triggered an acute nocifensive response and heat hyperalgesia. Selective pharmacological inhibition of TrkA and p75NTR dissociated these pain modalities: acute nociception and heat hyperalgesia were TrkA-dependent, while mechanical allodynia and cold hypersensitivity were mediated by p75NTR. Using adeno-associated viral (AAV) vectors and cell-specific Cre recombinase mouse lines targeting neurons or Schwann cells, we reported that acute nociception and heat hyperalgesia are exclusively neuronal, whereas mechanical allodynia and cold hypersensitivity are initiated and maintained by p75NTR activation in Schwann cells. This process involves reactive oxygen species (ROS) overproduction, which sustains itself *via* Schwann cell TRPA1 activation and concurrently activates neuronal TRPA1, thereby maintaining persistent mechanical and cold hypersensitivity.

Our findings redefine NGF signaling in pain by identifying a non-neuronal, Schwann cell-dependent p75NTR-ROS-TRPA1 axis as a key driver of chronic hypersensitivity. Targeting Schwann cell NGF signaling and redox-dependent intercellular crosstalk may open new therapeutic opportunities beyond neuron-centric strategies, providing a conceptual framework for the development of more selective and disease-modifying analgesic interventions.

## RESULTS

### TrkA mediates NGF-induced acute nociception and heat heyperalgesia, whereas p75NTR drives mechanical allodynia and cold hypersensitivity

NGF release, which frequently occurs at sites of peripheral tissue injury, can rapidly activate peripheral nociceptor terminals, leading to acute nociception, or promote the development of persistent pain states manifested as mechanical hyperalgesia and/or allodynia (Khodorova *et al*, 2013; Pezet & McMahon, 2006). In C57BL/6J mice, intraplantar injection of NGF elicited a dose-dependent (0.1, 0.5, 1 and 5 µg/paw) acute nociceptive response together with a dose- and time-dependent (6 hours) mechanical allodynia (Fig. 1a,b). Previous studies have shown that intradermal administration of NGF induces local thermal hypersensitivity (Dyck *et al*, 1997; Eskander *et al*, 2015). Consistently, we observed that intraplantar NGF produced long-lasting hypersensitivity to both heat and cold stimuli (Fig. 1b). Since both the acute nociceptive response and mechanical allodynia as well as thermal sensitivity were unchanged in male and female mice, all subsequent experiments were conducted in male mice (Suppl. Fig. 1a).

**Figure 1.**
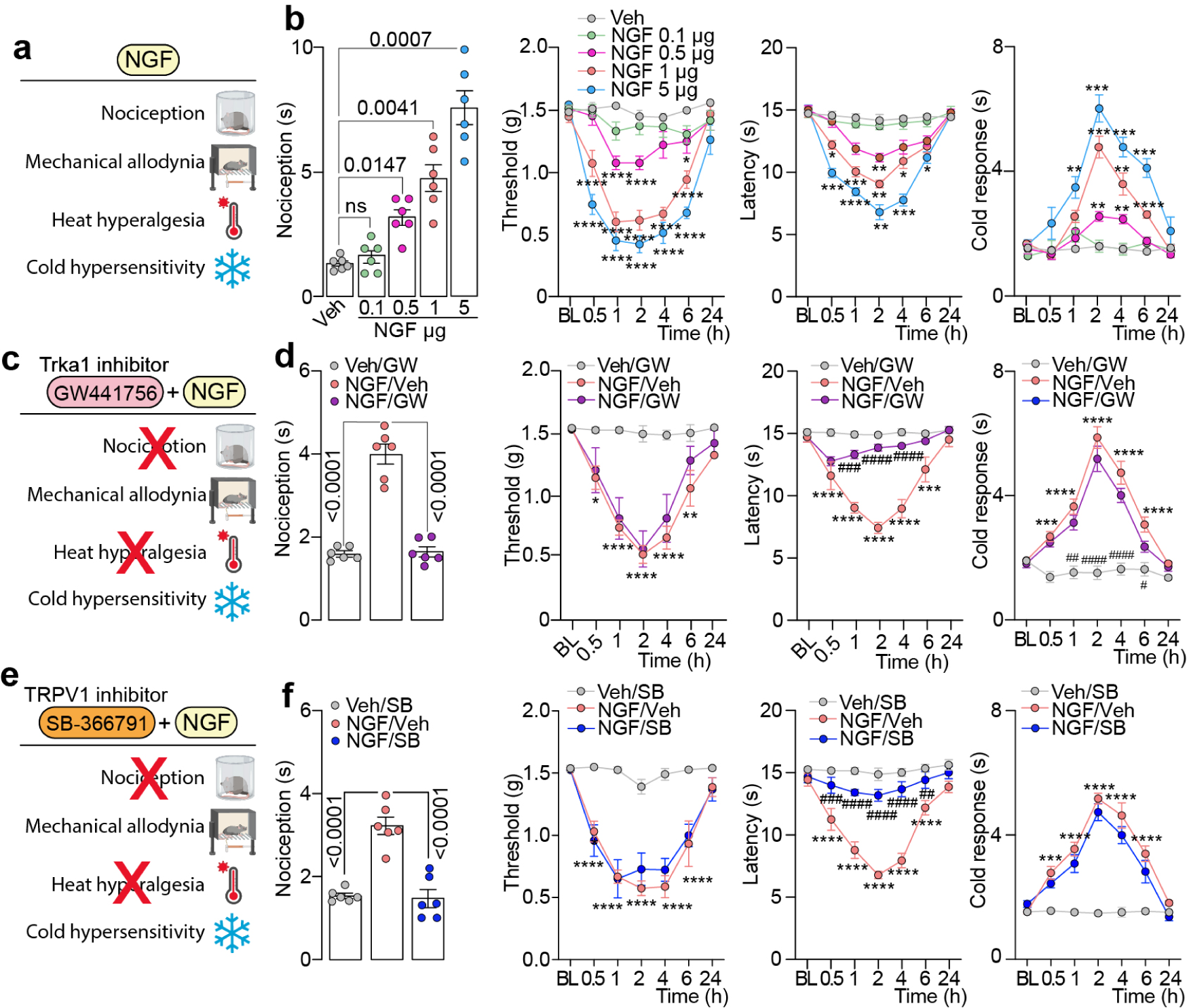
NGF-induced pain behaviors in mice and the contribution of TrkA and TRPV1 signaling. **(a)** Schematic representation of the pain behaviors (acute nociception, mechanical allodynia, heat hyperalgesia, and cold hypersensitivity) evaluated following intraplantar (i.pl./10 µl) NGF administration. **(b)** Dose-dependent acute nociception and dose- and time-dependent mechanical allodynia, heat hyperalgesia and cold hypersensitivity after intraplantar (i.pl./10 µl) injection of NGF (0.1, 0.5, 1 and 5 µg/paw) or vehicle (Veh) in C57BL/6J mice. **(c)** Schematic representation of pharmacological inhibition of TrkA signaling by GW441756 in NGF-induced pain behaviors. **(d)** acute nociception, mechanical allodynia, heat hyperalgesia and cold hypersensitivity after i.pl. NGF (1 µg) or Veh in C57BL/6J pretreated with GW441756 (GW, 10 nmol) or Veh. **(e)** Schematic representation of pharmacological inhibition of TRPV1 by SB-366791 in NGF-induced pain behaviors. **(f)** acute nociception, mechanical allodynia, heat hyperalgesia and cold hypersensitivity after i.pl. NGF (1 µg) or Veh in C57BL/6J pretreated with SB-366791 (SB, 10 nmol) or Veh. (n=6 mice per group). Data are mean ± s.e.m. 1-way or 2-way ANOVA, Bonferroni correction. *P<0.05, **P<0.01, ***P<0.001, ****P<0.0001 vs. Veh ^##^P<0.01, ^####^P<0.0001 vs. NGF/Veh.

NGF signals through two main receptors: the p75NTR and the TrkA (Minnone *et al*., 2017). The mature form of NGF exhibits a higher binding affinity for TrkA, while retaining the ability to engage p75NTR; in contrast, its precursor proNGF binds preferentially p75NTR (Mantyh *et al*, 2011; Mizumura & Murase, 2015). TrkA is highly expressed in nociceptive neurons (Fang *et al*, 2005). Accordingly, pretreatment with a selective TrkA inhibitor (GW441756) prevented NGF-induced nociception and heat hyperalgesia (Fig. 1 c,d). Recent evidence indicates that NGF can bind TrkA and Neuropilin-1 (NRP1), forming a ternary complex capable of activating TRPV1 and increasing nociceptor excitability (Peach *et al*., 2024). In line with this, we found that pretreatment with the TRPV1 selective inhibitor SB-366791 reduced NGF-induced nociception and heat hyperalgesia (Fig. 1e,f). Notably, inhibition of either TrkA or TRPV1 did not affect NGF-induced mechanical allodynia or cold hypersensitivity (Fig. 1c-f).

We also reported that the intraplantar injection of cleavage resistant proNGF evoked a dose-dependent, long-lasting mechanical allodynia, as well as cold hypersensitivity but not heat hyperalgesia (Fig. 2a,b). ProNGF also failed to evoke an acute nociceptive response (Fig. 2a,b). Same results were obtained in female mice (Suppl. Fig. 1b).

**Figure 2.**
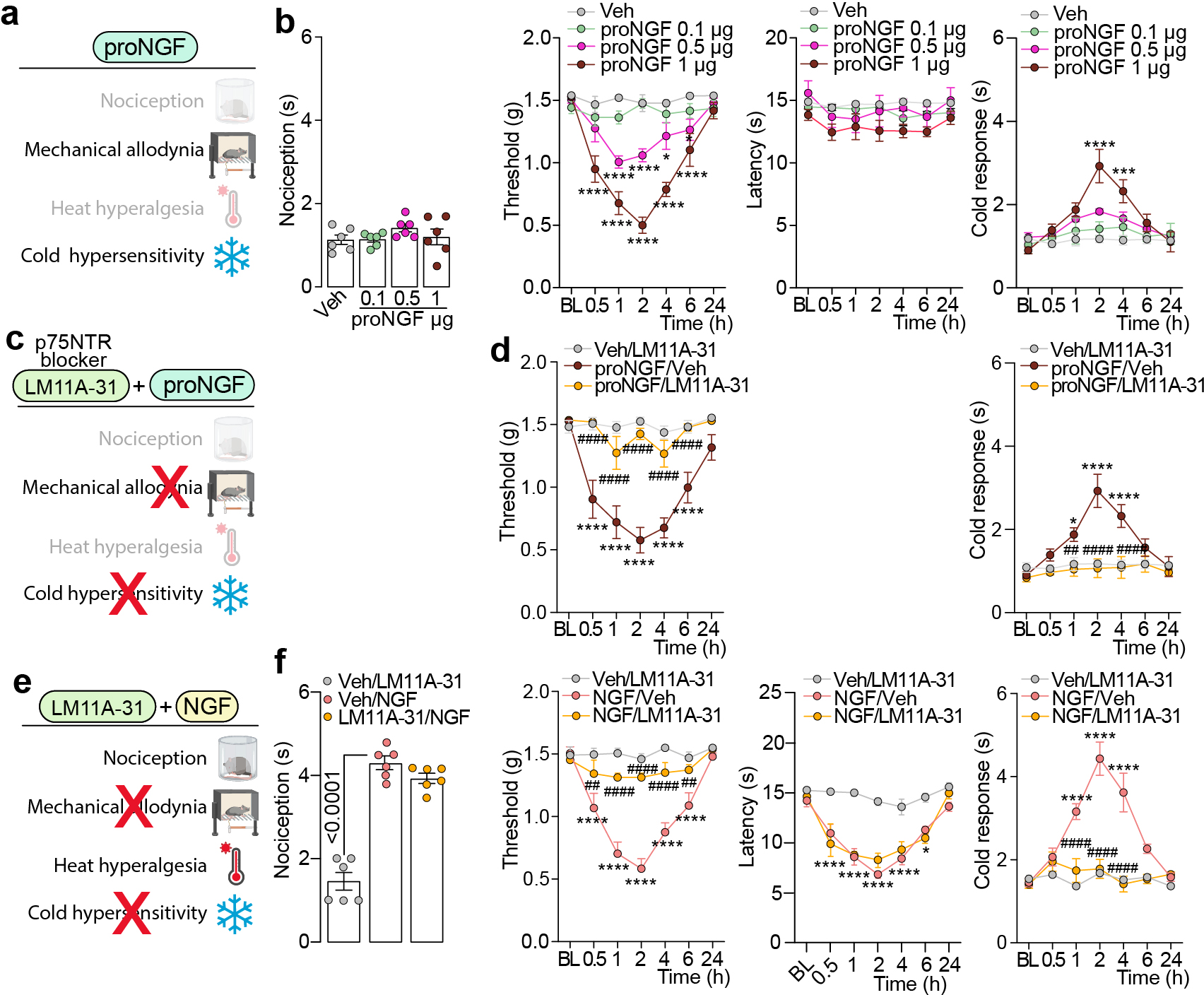
NGF- and proNGF-induced pain behaviors in mice and the contribution of p75NTR signaling. **(a)** Schematic representation of the pain behaviors (acute nociception, mechanical allodynia, heat hyperalgesia, and cold hypersensitivity) evaluated following intraplantar (i.pl./10 µl) proNGF administration. **(b)** Dose-dependent acute nociception and dose- and time-dependent mechanical allodynia, heat hyperalgesia and cold hypersensitivity after proNGF (0.1, 0.5, 1 µg/10 µl, i.pl.) or vehicle (Veh) in C57BL/6J mice. Schematic representation of LM11A-31 targeting p75NTR receptor in **(c)** proNGF and **(e)** NGF-induced pain behaviors. **(d)** mechanical allodynia and cold hypersensitivity after i.pl. proNGF (1 µg) or Veh in C57BL/6J pretreated with LM11A-31 (100 nmol) or Veh. **(f)** acute nociception, mechanical allodynia, heat hyperalgesia and cold hypersensitivity after i.pl. NGF (1 µg) or Veh in C57BL/6J pretreated with LM11A-31 (100 nmol) or Veh. (n=6 mice per group). Data are mean ± s.e.m. 1-way or 2-way ANOVA, Bonferroni correction. *P<0.05, ***P<0.001, ****P<0.0001 vs. Veh, ^##^P<0.01, ^####^P<0.0001 vs. NGF/Veh, proNGF/Veh.

We then used LM11A-31, a non-peptide ligand of p75NTR to assess its role in NGF and proNGF induced pain. Local (intraplantar) pretreatment with LM11A-31 significantly prevented in C57BL/6J mice proNGF- and NGF-dependent mechanical allodynia and cold hypersensitivity, while leaving unaffected NGF-induced acute nociception and heat hyperalgesia (Fig 2c-f). Overall, these data suggest that acute nociception and heat hypersensitivity are preferentially mediated by TrkA activation and TRPV1 sensitization, whereas mechanical allodynia and cold hypersensitivity are selectively driven by p75NTR signaling.

### Schwann cell p75NTR activation modulates mechanical allodynia and cold hypersensitivity

Given our previous evidence identifying Schwann cells as active modulators of chronic pain(De Logu *et al*, 2021; De Logu *et al*., 2022; Landini *et al*., 2023; Nassini *et al*., 2025; Titiz *et al*., 2024), we next investigated whether p75NTR signaling in Schwann cells mediates NGF-induced hypersensitivity. Beyond its expression in nociceptive neurons, p75NTR is highly expressed in both mature and immature Schwann cells, and results upregulated following nerve injury(Cosgaya *et al*, 2002; Zhou & Li, 2007). Based on this expression pattern, we wondered whether the activation of Schwann cell p75NTR preferentially contributes to NGF dependent pain.

To specifically define the contribution of Schwann cell p75NTR to NGF- and proNGF-induced mechanical allodynia and cold hypersensitivity, we selectively silenced p75NTR in Schwann cells. We employed a previously validated (Titiz *et al*., 2024) Cre-dependent AAVrh10 vector carrying a *loxP* flanked shRNA for Schwann cell selective *Ngfr* silencing (shRNA-*Ngfr*) in combination with a *Plp^Cre^* driver that functions as a lineage tracer to express shRNA-*Ngfr* only in Schwann cells of *Plp^Cre^* mice. In *Plp-Ngfr* intraplantar injection of NGF or proNGF resulted in a significant reduction of mechanical allodynia and cold hypersensitivity, compared to control mice (Fig. 3a-d). Conversely, acute nociception or heat hypersensitivity mediated by NGF were not affected (Fig. 3a,b).

**Figure 3.**
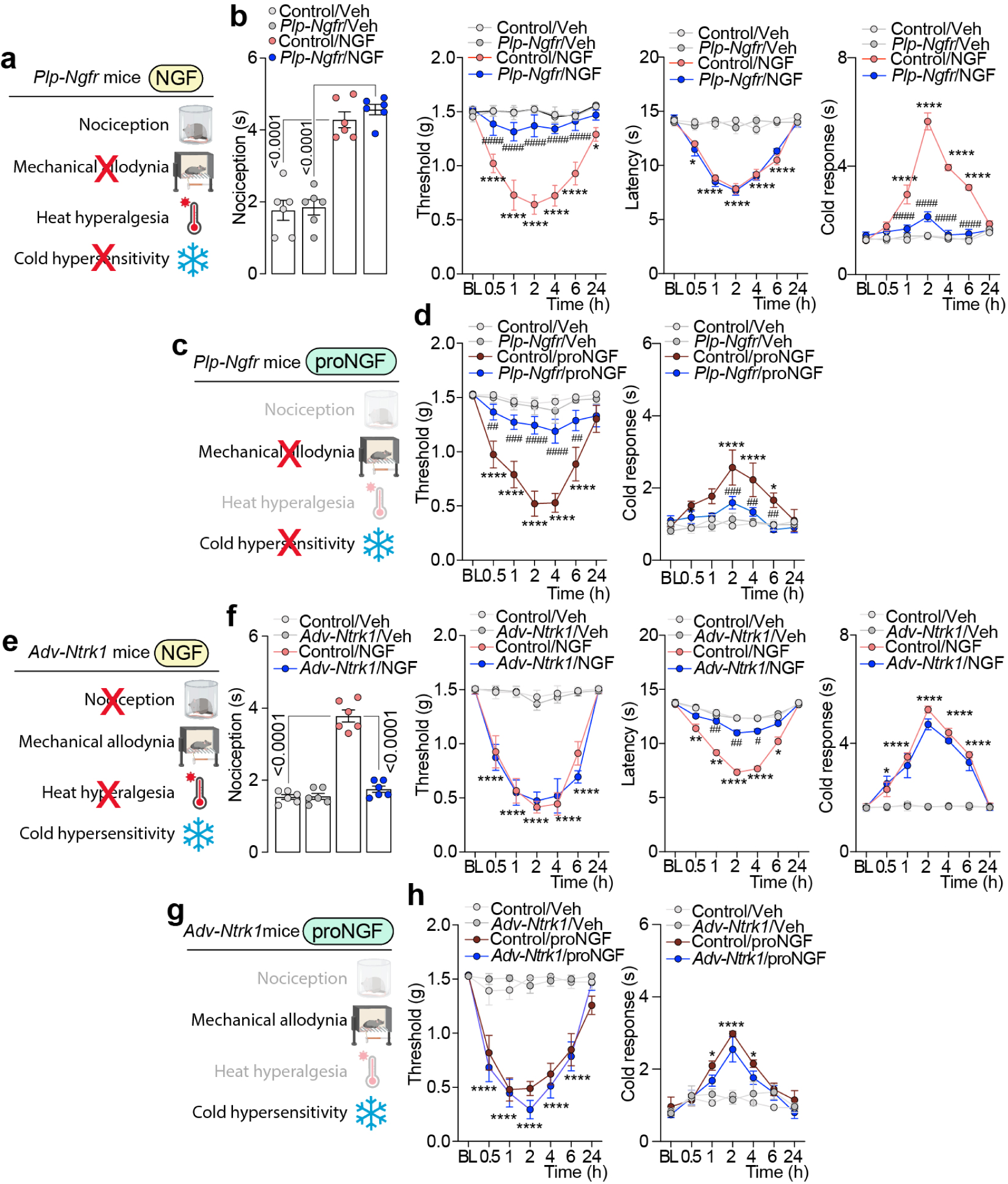
Distinct role of NGF and proNGF signaling in Schwann cells and sensory neurons in pain behaviors in mice. **(a)** Schematic representation of NGF-induced pain behaviors (acute nociception, mechanical allodynia, heat hyperalgesia, and cold hypersensitivity) *in P1p-Cre* mice infected with AAV for selective silencing of p75NTR (*Plp-Ngfr* mice) in Schwann cells. **(b)** acute nociception, mechanical allodynia, heat hyperalgesia and cold hypersensitivity after intraplantar (i.pl./10 µl) injection of NGF (1 µg) or vehicle (Veh) in *Plp-Ngfr* or Control mice. **(c)** Schematic representation of proNGF-induced pain behaviors in *Plp-Ngfr* mice. **(d)** mechanical allodynia and cold hypersensitivity after i.pl. NGF (1 µg) or Veh in *Plp-Ngfr* or Control mice. (**e)** Schematic representation of NGF-induced pain behaviors in *Adv-Cre* mice infected with AAV for selective silencing of TrkA (*Adv- Ntrk1* mice) in primary sensory neurons. **(f)** acute nociception, mechanical allodynia, heat hyperalgesia and cold hypersensitivity after after i.pl. NGF (1 µg) or Veh in *Adv-Ntrk1* or Control mice. **(g)** Schematic representation of proNGF-induced pain behaviors in *Adv Ntrk1* mice. **(h)** mechanical allodynia and cold hypersensitivity after i.pl. proNGF (1 µg) or Veh in *Adv-Ntrk1* or Control mice. (n=6 mice per group). Data are mean ± s.e.m. 1-way or 2-way ANOVA, Bonferroni correction. *P<0.05, **P<0.01, ****P<0.0001 vs. Control/Veh ^##^P<0.01, ^####^P<0.0001 vs. Control/NGF and Control/proNGF.

To confirm the predominant role of primary sensory neurons in NGF- and proNGF dependent acute nociception and thermal heat hypersensitivity we induced the selective silencing of TrkA in primary sensory neurons in *Adv^Cre^* mice. We used a previously validated (Chieca *et al*, 2026) Cre- dependent AAVPHP.S with *loxP* flanked shRNA for primary sensory neurons selective *Ntrk1* silencing (shRNA-*Ntrk1*) and a *Adv^Cre^* to express shRNA-*Ntrk1* only in primary sensory neurons in *Adv^Cre^* mice. *Adv*-*Ntrk1* displayed markedly reduced nociceptive responses and heat hypersensitivity following NGF injection (Fig. 3e,f), whereas mechanical allodynia and cold hypersensitivity induced by both NGF and proNGF were unchanged (Fig. 3e-h). These findings demonstrate a functional segregation between Schwann cells and sensory neurons in NGF-dependent pain processing.

### p75NTR activation induces early calcium mobilization and mitochondrial ROS production

Schwann cells, through their broad repertoire of receptors and enzymes, can sense algogenic stimuli and activate intracellular signaling pathways that result in the overproduction of reactive oxygen species (ROS). Through the involvement and activation of the transient potential ankyrin 1 (TRPA1) channel, the ROS overproduction establishes a feed forward mechanism that promotes and maintains the chronic pain (De Logu *et al*, 2019; De Logu *et al*., 2022; De Logu *et al*, 2017). The p75NTR receptor signaling is also closely associated with oxidative stress: ROS can stimulate p75NTR endocytosis, thereby increasing its susceptibility to metalloprotease-mediated cleavage (Pokharel *et al*, 2024), while p75NTR activation itself has been shown to induce the expression of regulatory subunits of NADPH oxidase, promoting ROS generation (Pensabene *et al*, 2025).

First, to investigate whether NGF- and proNGF induced mechanical allodynia and cold hypersensitivity were dependent on ROS production, we administered the antioxidant N-tert-butyl- α-phenylnitrone (PBN). Pretreatment of C57BL/6J mice with PBN significantly attenuated the development of mechanical allodynia and cold hypersensitivity by NGF and proNGF, without affecting acute nociception or heat hypersensitivity (Fig. 4a-d).

**Figure 4.**
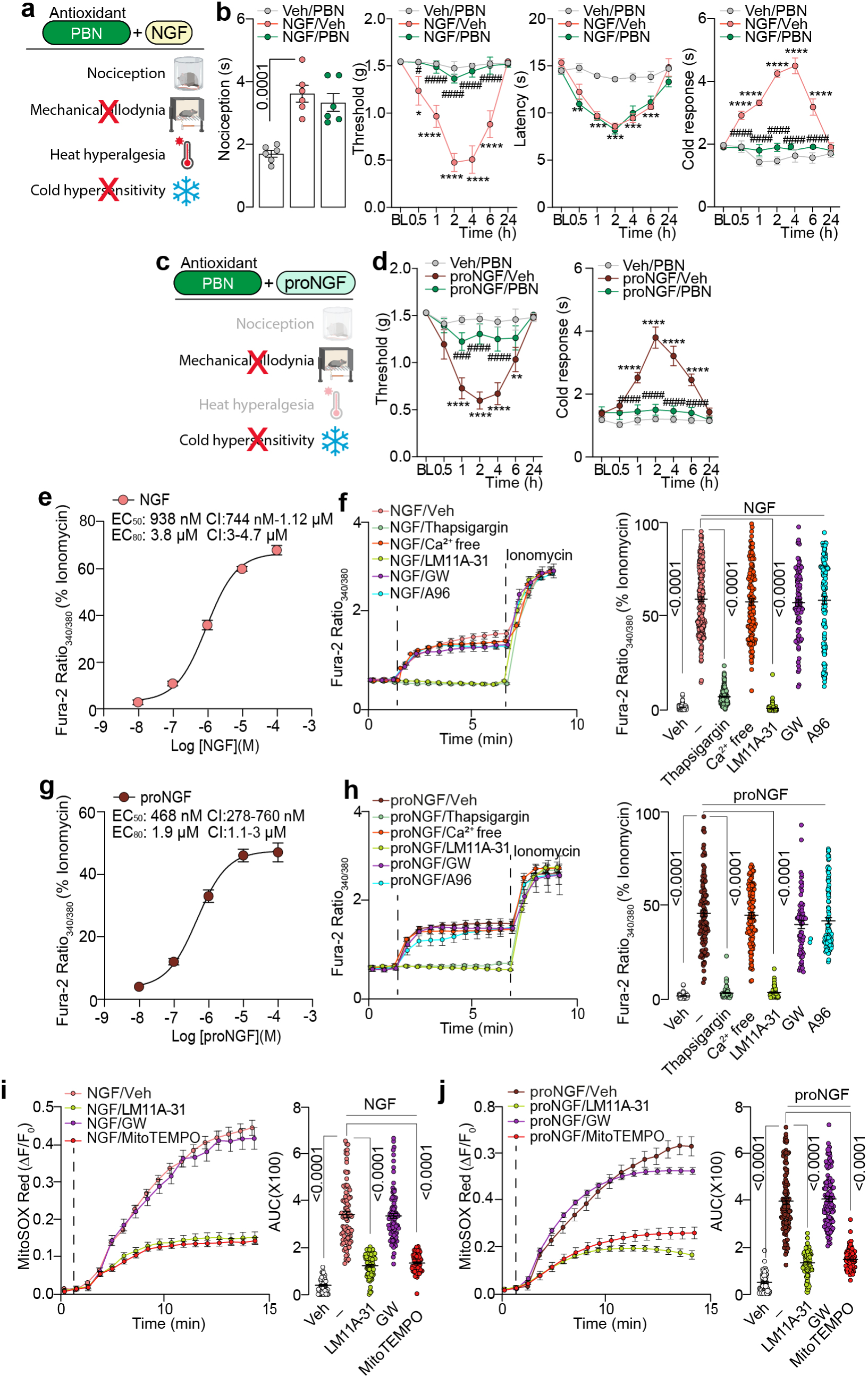
NGF- and proNGF-induced p75NTR activation promotes mitochondrial ROS production in Schwann cells, mechanical allodynia, and cold hypersensitivity. **(a)**Schematic representation of ROS inhibition by N-tert-butyl-α-phenylnitrone (PBN) in NGF-induced pain behaviors (acute nociception, mechanical allodynia, heat hyperalgesia, and cold hypersensitivity). **(b)** acute nociception, mechanical allodynia, heat hyperalgesia and cold hypersensitivity after intraplantar (i.pl./10 µl) NGF (1 µg) or vehicle (Veh) in C57BL/6J mice pretreated with PBN (670 nmol) or Veh. (**c)** Schematic representation of ROS inhibition by PBN in proNGF-induced pain behaviors. **(d)** Mechanical allodynia and cold hypersensitivity after i.pl. proNGF (1 µg) or Veh in C57BL/6J mice pretreated with PBN (670 nmol) or Veh. (n=6 mice per group). Concentration response curve of Ca^2+^ evoked by **(e)** NGF or **(g)** proNGF in human Schwann Cells (SCs) (cells number: NGF: 10 nM=238, 100 nM=179, 1µM=203, 10µM=196, 100µM=127; proNGF: 10 nM=77, 100 nM=87, 1µM=92, 10µM=106, 100µM=87). Typical traces and pooled data of Ca^2+^ responses in SCs after **(f)** NGF (4 µM) or **(h)** proNGF (2 µM) in presence of thapsigargin (2 µM), Ca^2+^ free medium, LM11A-31 (1 µM), GW441756 (GW, 1 µM), A-967079 (A96, 30 µM) or Veh (0.001% DMSO) (cells number: NGF: Veh=131, NGF/Veh=196, NGF/Thapsigargin=159, NGF/ Ca^2+^ free=122, NGF/ LM11A-31 =99, NGF/GW=93, NGF/A96=127; proNGF: Veh=103, proNGF/Veh=106, proNGF/Thapsigargin=104, proNGF/Ca^2+^free=100, proNGF/LM11A-31=76, proNGF/GW441756=64, proNGF/A96=99). **(i)** Typical traces and cumulative data (area under the curve, AUC) of MitoSOX Red fluorescence in SCs stimulated with **(i)** NGF (4 µM) or **(j)** proNGF (2 µM) in the presence of LM11A-31 (1 µM), GW (1 µM), Mito TEMPO (5 µM) or Veh (0.001% DMSO) (cells number: NGF: Veh=74, NGF/Veh=88, NGF/LM=76, NGF/GW=87, NGF/MitoTEMPO=80; proNGF: Veh=70, proNGF/Veh=107, proNGF/LM=75, proNGF/GW=105, proNGF/MitoTEMPO=75). Data are mean ± s.e.m. 1-way or 2-way ANOVA, Bonferroni correction. *P<0.05, **P<0.01, ***P<0.001, ****P<0.0001 vs. Veh/PBN ^###^P<0.001, ^####^P<0.0001 vs. NGF/Veh and proNGF/Veh.

We then assessed whether NGF and proNGF were able to increase intracellular calcium levels in Schwann cells. Stimulation with either NGF or proNGF induced a rapid increase in intracellular calcium (Fig. 4e-h). This early calcium response was abolished by preincubation of Schwann cells with thapsigargin but was not affected by calcium-free extracellular medium, indicating that it mainly resulted from mobilization of intracellular stores rather than extracellular calcium influx (Fig. 4f,h). Consistently, the rapid calcium increase was prevented by the p75NTR inhibitor (LM11A-31), but not by TrkA inhibitior (GW441756) (Fig. 4f,h). These findings are in line with previous evidence showing that activation can promote the increase in calcium through mobilization of intracellular calcium stores (De Bernardi *et al*, 1996). Importantly, the kinetics of this early calcium rise were too rapid to be compatible with a TRPA1-dependent mechanism driven by newly generated ROS. In agreement with this interpretation, the selective TRPA1 antagonist (A967079) did not prevent the initial calcium increase (Fig. 4f,h). Together, these findings indicate that the early calcium transient is primarily mediated by p75NTR -dependent release of calcium from intracellular stores, rather than by TRPA1-mediated calcium entry.

Previous studies reported that p75NTR activation can induce superoxide production through mitochondrial pathways involving superoxide dismutase 1 (SOD1)(Pehar *et al*, 2007). Therefore, we next investigated whether the early calcium response was associated with mitochondrial ROS generation. Using the MitoSOX Red probe, we observed that stimulation with NGF or proNGF markedly increased mitochondrial superoxide production in Schwann cells (Fig. 4i,j). This response was prevented by p75NTR inhibition (LM11A-31), but not by TrkA inhibition (GW441756) (Fig. 4i,j). In addition, pretreatment with the mitochondria-targeted antioxidant MitoTEMPO abolished the increase in MitoSOX fluorescence induced by NGF or proNGF, supporting the involvement of mitochondrial superoxide downstream of p75NTR activation (Fig. 4i,j). Collectively, these findings suggest that early p75NTR-dependent calcium mobilization precedes and likely promotes mitochondrial ROS generation in Schwann cells.

### TRPA1 mediates delayed calcium influx and feed-forward oxidative signaling

The analysis of intracellular calcium dynamics revealed that NGF and proNGF induced a biphasic calcium response in Schwann cells. In addition to the rapid transient phase (from 0 to 5 minutes), we observed a delayed calcium increase with slower kinetics (from 5 to 20 minutes) (Fig. 5a,b). Unlike the early response, this delayed phase was markedly reduced by the TRPA1 antagonist A967079 and was markedly impaired in calcium-free extracellular medium, indicating that it depends, at least in part, on TRPA1-mediated extracellular calcium influx (Fig. 5a,b). The delayed kinetics of this response are consistent with the time required for p75NTR-dependent mitochondrial ROS production, which may subsequently activate TRPA1.

**Figure 5.**
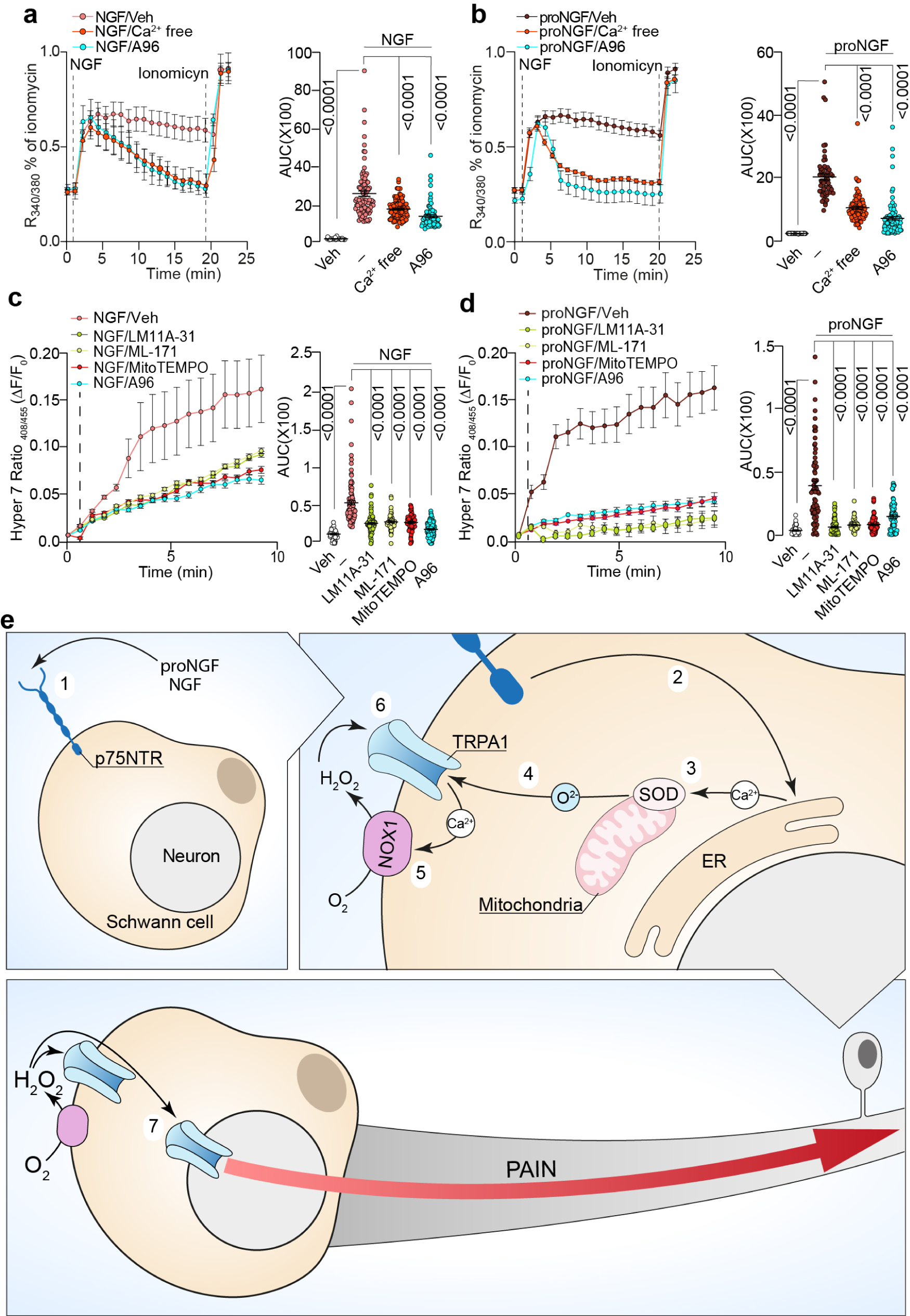
Schwann cell TRPA1 drives NGF and proNGF delayed intracellular calcium increase and oxidative stress signaling. Typical traces and pooled data of Ca^2+^ responses in human Schwann Cells (SCs) after **(a)** NGF (4 µM) or **(b)** proNGF (2 µM) in presence of Ca^2+^ free medium, A967079 (A96, 30µM) or vehicle (Veh, 0.001% DMSO) (cells number: NGF: Veh=34, NGF/Veh=91, NGF/Ca^2+^free=88, NGF/A96=70, proNGF: Veh=104, proNGF/Veh=62, proNGF/Ca^2+^ free=87, proNGF/A96=98). Typical traces and pooled data of Hyper 7 fluorescence after **(c)** NGF (4 µM) or **(d)** proNGF (2 µM) in presence of LM11A-31 (1 µM) ML-171 (1 µM), MitoTEMPO (5 µM), A96 (30 µM) or Veh (0.001% DMSO) (cells number: NGF: Veh=51, NGF/Veh=116, NGF/LM=94, NGF/ML-171=55, NGF/MitoTEMPO=75, NGF/A96=114; proNGF: Veh=40, proNGF/Veh=72, proNGF/ LM11A-31 =84, proNGF/ML-171=81, proNGF/MitoTEMPO=101, proNGF/A96=66). Data are mean ± s.e.m. 1-way ANOVA, Bonferroni correction. **(e)** Schematic representation of the signaling cascade induced by NGF- and proNGF-mediated p75NTR activation in Schwann cells. **(1)** NGF and proNGF activate p75NTR in Schwann cells, leading to **(2)** Ca²⁺ release from intracellular stores and **(3)** superoxide dismutase- (SOD)-dependent superoxide (O₂⁻) generation. **(4)** Reactive oxygen species activate TRPA1, resulting in extracellular Ca²⁺ influx and **(5)** Ca²⁺-dependent activation of NADPH oxidase 1 (NOX1). **(6)** NOX1-derived hydrogen peroxide (H₂O₂) sustains a feed-forward mechanism by further activating TRPA1 in Schwann cells and, simultaneously, (**7)** activates neuronal TRPA1 to signal pain.

Thus, the most parsimonious interpretation of our findings is that NGF and proNGF-induced activation of p75NTR first triggers a rapid release of calcium from intracellular stores that promotes mitochondrial superoxide production. Superoxide then activates TRPA1 with delayed kinetics, leading to extracellular calcium entry. This TRPA1-dependaaent calcium influx may in turn activate NOX1, promoting H₂O₂ release and establishing a feed-forward oxidative mechanism that sustains TRPA1 activation and amplifies the pro-nociceptive signal in Schwann cells.

To directly test the proposed p75NTR/SOD1/TRPA1/NOX1/H₂O₂ axis, we measured H₂O₂ production in Schwann cells using the genetically encoded fluorescent probe HyPer7. Stimulation with either NGF or proNGF induced a delayed increase in HyPer7 fluorescence, with slower kinetics than those observed for mitochondrial superoxide production (Fig. 5c,d). This temporal delay is consistent with the interpretation that H₂O₂ generation occurs downstream of the initial p75NTR- dependent mitochondrial superoxide signal. Importantly, the NGF- and proNGF-induced increase in HyPer7 fluorescence was prevented by pretreatment with a p75NTR inhibitor (LM11A-31), the mitochondria-targeted antioxidant (MitoTEMPO), the TRPA1 antagonist (A967079), and the NOX1 inhibitor (ML171) (Fig. 5c,d). These pharmacological data support a sequential mechanism in which p75NTR activation promotes mitochondrial superoxide production, which subsequently activates TRPA1. TRPA1-dependent calcium influx then engages NOX1 activity, leading to delayed H₂O₂ generation (Fig. 5e).

To specifically assess the *in vivo* relevance of TRPA1, we used the selective TRPA1 antagonist A967079. Pharmacological inhibition of TRPA1 significantly reduced NGF- and proNGF-induced mechanical allodynia and cold hypersensitivity without affecting acute nociceptive responses or heat hyperalgesia (Fig. 6a-d). Similarly, selective silencing of *Trpa1* in Schwann cells (*Plp-Trpa1^fl/fl^* mice), markedly attenuated NGF- and proNGF-induced mechanical allodynia and cold hypersensitivity, while preserving nociceptive and heat responses (Fig. 6e-h). Consistently, sciatic nerve homogenates from *Plp-Trpa1^fl/fl^* mice displayed significantly lower ROS levels after NGF or proNGF administration compared with controls (Fig. 6i).

**Figure 6.**
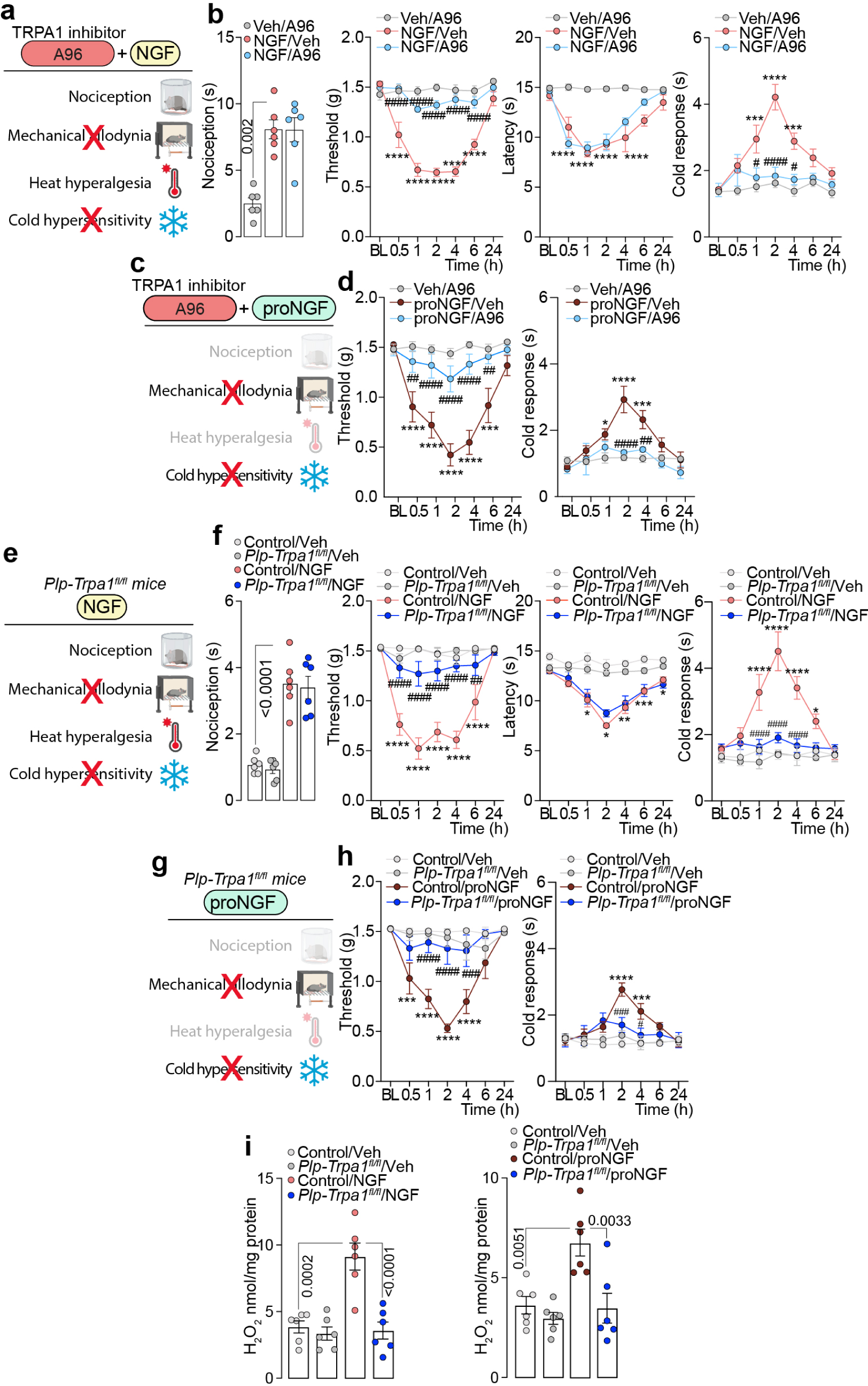
Schwann cell TRPA1 drives NGF and proNGF dependent mechanical allodynia and cold hypersensitivity. **(a)** Schematic representation of TRPA1 inhibition by A-967079 in NGF-induced pain behaviors (acute nociception, mechanical allodynia, heat hyperalgesia, and cold hypersensitivity). **(b)** acute nociception, mechanical allodynia, heat hyperalgesia and cold hypersensitivity after intraplantar (i.pl./10 µl) NGF (1 µg) or vehicle (Veh) in C57BL/6J mice pretreated with A-967079 (A96, 300 nmol, i.pl.) or Veh. **(c)** Schematic representation of TRPA1 inhibition by A-967079 in proNGF- induced pain behaviors. **(d)** mechanical allodynia and cold hypersensitivity after i.pl. proNGF (1 µg) or Veh in C57BL/6J mice pretreated with A96 (300 nmol, i.pl.) or Veh. **(e)** Schematic representation of NGF-induced pain behaviors *in P1p-Trpa1^fl/fl^* mice. **(f)** acute nociception, mechanical allodynia, heat hyperalgesia and cold hypersensitivity after i.pl. NGF (1 µg) or Veh in *in P1p-Trpa1^fl/fl^* or Control mice. **(g)** Schematic representation of proNGF-induced pain behaviors *in P1p-Trpa1^fl/fl^* mice. **(h)** mechanical allodynia and cold hypersensitivity after i.pl. proNGF (1 µg) or Veh in *in P1p-Trpa1^fl/fl^* or control mice. (**i,j**) H_2_O_2_ content in sciatic nerve tissue homogenates of *P1p-Trpa1^fl/fl^* or Control mice after i.pl. NGF (1 µg) or veh **(i)** and proNGF (1 µg) or Veh **(j).** (n=6 mice per group). Data are mean ± s.e.m. 1-way or 2-way ANOVA, Bonferroni correction. *P<0.05, ***P<0.001, ****P<0.0001 vs. Veh/A96, Control/Veh, ^#^P<0.05, ^##^P<0.01, ^###^P<0.001, ^####^P<0.0001 vs NGF/Veh, ProNGF/Veh, Control/NGF, Control/ProNGF.

Collectively, these findings identify a temporally defined p75NTR-dependent oxidative signaling cascade in Schwann cells, in which early intracellular calcium mobilization and mitochondrial ROS production drive delayed TRPA1 activation and feed-forward oxidative amplification, selectively sustaining mechanical allodynia and cold hypersensitivity.

## Discussion

Here, we reveal distinct cellular and molecular mechanisms underlying the different pain phenotypes driven by NGF and proNGF. We show that acute nociception and heat hyperalgesia are primarily mediated by TrkA-TRPV1 signaling in primary sensory neurons, whereas mechanical allodynia and cold hypersensitivity are selectively driven by p75NTR-dependent activation of Schwann cells. NGF has long been recognized as a central mediator of inflammatory and neuropathic pain through its actions on TrkA-expressing nociceptors (Lewin & Mendell, 1993; Mantyh *et al*., 2011; Pezet & McMahon, 2006; Woolf *et al*., 1994). By sensitizing TRPV1 channels, enhancing neuronal excitability, and promoting both peripheral and central sensitization, NGF contributes to the development and maintenance of thermal hyperalgesia and spontaneous pain (Amaya *et al*, 2004; Chuang *et al*, 2001; Mills *et al*, 2013; Stein *et al*, 2006). To date, experimental evidence regarding the mechanisms underlying NGF-induced mechanical allodynia and, in particular, thermal hypersensitivity remains partly conflicting. Previous studies have shown that mechanical allodynia and heat hyperalgesia induced by repeated NGF administration are attenuated by pretreatment with the TRPV1 antagonist capsazepine (Eskander *et al*., 2015). In contrast, NGF-induced heat hyperalgesia has also been reported to be predominantly inhibited by intraplantar administration of an anti-p75NTR antibody (Watanabe *et al*, 2008), suggesting a contribution of p75NTR signaling to this response. Further complexity arises from studies using genetically modified NGF or p75NTR- deficient models. The NGF variant associated with hereditary sensory and autonomic neuropathy type V, characterized by the R100W substitution in the mature NGF sequence (NGF^R100W), shows markedly reduced binding and activation of p75NTR. This mutant failed to induce thermal hyperalgesia while retaining the ability to evoke persistent mechanical hypersensitivity (Sung *et al*, 2018). Conversely, NGF administration in p75NTR-deficient mice did not reveal major differences in either mechanical or heat hyperalgesia compared with wild-type animals (Bergmann *et al*, 1998). In addition, treatment with a polyclonal anti-p75NTR antibody was reported not to prevent NGF-induced thermal hyperalgesia, but rather to shorten its duration (Khodorova *et al*, 2017). Our study addresses these apparently divergent findings by dissecting the cellular contribution of p75NTR signaling to distinct NGF- and proNGF-induced pain modalities, including mechanical allodynia, acute nociception, and heat and cold hypersensitivity. Our findings identify Schwann cells as a critical cellular site of p75NTR signaling. Indeed, Schwann cell-selective silencing of p75NTR markedly attenuated NGF- and proNGF-induced mechanical and cold hypersensitivity, while leaving acute nociception and heat hyperalgesia largely unaffected. These results therefore provide a cellular framework that may help reconcile previous apparently conflicting observations and define a Schwann cell-dependent component of p75NTR-mediated NGF signaling. More broadly, they support a functional segregation between predominantly neuronal TrkA-dependent signaling and Schwann cell p75NTR-dependent mechanisms in the modulation of distinct pain modalities.

Notably, NGF and proNGF-induced Schwann activation involve a biphasic calcium response, mitochondrial ROS generation downstream to p75NTR activation, and delayed TRPA1-dependent oxidative amplification. These findings are consistent with previous evidence linking p75NTR signaling to oxidative stress, mitochondrial dysfunction, and NADPH oxidase activation (Ferrari *et al*, 2010; Pokharel *et al*., 2024; Tammariello *et al*, 2000). Importantly, our results extend these observations by defining the temporal organization of calcium and ROS signaling downstream of p75NTR activation. The early calcium store mobilization occurred rapidly after stimulation, and was abolished by p75NTR inhibition or thapsigargin treatment, but not by extracellular calcium removal or TRPA1 inhibition. Previous studies have reported that p75NTR activation can modulate intracellular calcium signaling through the release of calcium from endoplasmic reticulum stores, supporting our observations (De Bernardi *et al*., 1996). In contrast, the delayed calcium phase displayed slower kinetics and was markedly reduced by extracellular calcium removal or TRPA1 inhibition. The temporal delay between mitochondrial ROS generation and late calcium entry strongly supports a sequential mechanism in which p75NTR activation first induces mitochondrial superoxide production, which subsequently activates TRPA1. TRPA1 activation then promotes extracellular calcium influx, potentially engaging NOX1 activity and leading to secondary H₂O₂ production. Consistent with this model, HyPer7 imaging revealed delayed H₂O₂ generation following NGF or proNGF stimulation, and this response was prevented by inhibition of p75NTR, mitochondrial ROS, TRPA1, or NOX1. Together, these data support the existence of a p75NTR/mitochondrial ROS/TRPA1/NOX1 feed-forward pathway that amplifies oxidative signaling in Schwann cells.

TRPA1 has emerged as a critical sensor of oxidative stress and electrophilic mediators in chronic pain (Andersson *et al*, 2008; Bessac *et al*, 2008; Trevisani *et al*, 2007). ROS and lipid peroxidation products can directly activate TRPA1, thereby increasing neuronal excitability and neurogenic inflammation (Materazzi *et al*, 2008). Previous studies from our group demonstrated that Schwann cell TRPA1 sustains chronic pain through ROS-dependent feed-forward signaling (De Logu *et al*., 2019; De Logu *et al*., 2022; De Logu *et al*., 2017). The present findings extend this concept by identifying p75NTR as an upstream trigger of Schwann cell oxidative signaling. Importantly, pharmacological inhibition or selective Schwann cell deletion of TRPA1 significantly reduced NGF- and proNGF-induced mechanical and cold hypersensitivity, while preserving acute nociception and heat responses. These results indicate that Schwann cell TRPA1 selectively contributes to persistent hypersensitivity rather than acute nociceptive transmission.

The functional segregation between Schwann cells and sensory neurons highlighted in this study underscores the complexity of peripheral pain processing. While primary sensory neurons remain the principal transducers of noxious stimuli, Schwann cells are more widely emerging e as active modulators capable of selectively shaping specific pain modalities through oxidative and inflammatory signaling. These findings are consistent with growing evidence implicating peripheral glial cells in neuroimmune interactions and chronic pain maintenance (De Logu *et al*, 2020; Donnelly *et al*, 2020; Ji *et al*, 2018). This observation may also have important translational implications. Although NGF-neutralizing antibodies demonstrated efficacy in osteoarthritis and chronic pain, their clinical development has been limited by adverse effects and safety concerns (Lane *et al*, 2010; Miller *et al*, 2017). Our data suggests that selectively targeting downstream pathways, such as Schwann cell p75NTR signaling or ROS-TRPA1 activation, may provide more refined therapeutic strategies capable of reducing persistent hypersensitivity while preserving protective nociceptive functions. Moreover, the preferential involvement of proNGF in persistent pain raises the possibility that limiting proNGF accumulation or extracellular processing could represent an additional therapeutic opportunity.

In conclusion, our study identifies a dual and cell-specific mechanism of NGF- and proNGF-induced pain. Acute nociception and heat hyperalgesia are mediated by neuronal TrkA-TRPV1 signaling, whereas persistent mechanical and cold hypersensitivity depend on Schwann cell p75NTR activation and ROS-TRPA1 feed-forward signaling. However, future studies should explore how Schwann cell p75NTR signaling interacts with immune mediators, extracellular matrix remodeling, and axonal transport to shape chronic pain trajectories. It will also be important to determine whether similar mechanisms operate in human tissues and clinical pain conditions, particularly in inflammatory, cancer-related, and neuropathic syndromes. Single-cell transcriptomic and proteomic approaches may further refine our understanding of Schwann cell heterogeneity and its relevance to pain modulation.

## Methods

### Reagents

Unless otherwise indicated, reagents were obtained from Merck Life Science SRL (Milan, Italy). Recombinant nerve growth factor (NGF, #N-130) and cleavage-resistant proNGF (#N-255) were from Alomone Labs (Jerusalem, Israel).

### Animals

Male and female mice C57BL/6 J (Charles River, RRID: IMSR_JAX:000664) were used throughout (25-30 g, 6-8 weeks old). To generate mice in which the *Trpa1* gene was conditionally silenced in Schwann cells, homozygous 129S-Trpa1^tm2Kykw/J^ (floxed Trpa1, *Trpa1^fl/fl^*, RRID:IMSR_JAX: 008649 Jackson Laboratory) were crossed with hemizygous B6.Cg-Tg(Plp1-CreERT)3Pop/J mice (*Plp- Cre^ERT^*, RRID: IMSR_JAX:005975 Jackson Laboratory) expressing a tamoxifen-inducible Cre in Schwann cells (Plp1, proteolipid protein myelin 1) (De Logu *et al*., 2017). The progeny (*Plp- Cre^ERT^;Trpa1^fl/fl^*) was genotyped using PCR for *Trpa1* and *Plp-Cre^ERT^.* Mice that were negative for *Plp-Cre^ERT^* (*Plp-Cre^ERT–^;Trpa1^fl/fl^*) were used as control. Both positive and negative mice for *Cre^ERT^* and homozygous floxed Trpa1 (*Plp-Trpa1* and control respectively) were treated with intraperitoneal (i.p.) 4-hydroxytamoxifen (4-OHT, 1 mg/100 μL in corn oil once a day consecutively for 3 days). Some *Plp-Cre^ERT^*^+^ or *Plp-Cre^ERT^*^-^ were treated with 4-OHT (i.p., 1 mg/100 μL in corn oil, once a day consecutively for 3 days) before the infection with AAV for selective silencing of the different genes in Schwann cells. Hemizygous Advillin-Cre mice (Adv-Cre^+^) or their control (Adv-Cre^-^) (De Logu *et al*., 2017) were also used for the infection with AAV for selective silencing of the different genes in primary sensory neurons.

The group size of n=8 mice for behavioral experiments was determined by sample size estimation using G Power [v3.1 (Faul *et al*, 2007)] to detect the size effect in a post-hoc test with type 1 and 2 error rates of 5% and 20%, respectively. Allocation concealment of mice into the vehicle(s) or treatment groups was performed using a randomization procedure (http://www.randomizer.org/). The assessors were blinded to the identity of the animals (genetic background) or allocation to treatment groups. None of the animals were excluded from the study. Mice were housed in a temperature- and humidity-controlled vivarium (12 h dark/light cycle, free access to food and water, 5 animals per cage). At least 1 h before behavioral experiments, mice were acclimatized to the experimental room and behavior was evaluated between 9:00 am and 5:00 pm. Animals were anesthetized with a mixture of ketamine and xylazine (90 mg/kg and 3 mg/kg, respectively, i.p.) and euthanized with inhaled CO2 plus 10-50% O2.

The research conducted complies with all relevant ethical regulations. Behavioral studies followed Animal Research: Reporting of In Vivo Experiments (ARRIVE) guidelines (McGrath & Lilley, 2015). Animal experiments and sample collections were carried out according to the European Union (EU) guidelines for animal care procedures and Italian legislation (DLgs 26/2014) application of the EU Directive 2010/63/EU. All animal studies were approved by the Animal Ethics Committee of the University of Florence and the Italian Ministry of Health (permits no. 765/2016-PR).

### Treatment protocols

Mice received intraplantar (i.pl., 10 µl/paw) injection of NGF (0.1, 0.5, 1 and 5 µg), proNGF (0.1, 0.5, and 1 µg) or vehicle (0.9% NaCl). The selective TrkA inhibitor, GW441756 (10 nmol), the selective TRPV1 inhibitor, SB-366791 (10 nmol), or the p75NTR inhibitor LM11A-31 (100 nmol), the ROS inhibitor N-tert-butyl-α-phenylnitrone (PBN, 670 nmol) the TRPA1 antagonist (A967079, 300 nmol) or their vehicle (4% DMSO, 4% tween 80 in NaCl 0.9%) were administered (i.pl., 10 µl) 30 minutes before NGF or proNGF.

### Behavioral experiments

#### Acute nociception

Immediately after i.pl. injection, mice were placed inside a plexiglass chamber, and acute nociception response was assessed for 10 min by measuring the time (sec) that the animal spent in lifting, biting, licking, shaking the injected paw.

#### Hindpaw mechanical allodynia

Mechanical sensitivity was evaluated using von Frey filaments of increasing stiffness (0.02-2 g) applied to the plantar surface of the hind paw according to the up-and-down paradigm(Chaplan *et al*, 1994). The 50% mechanical paw-withdrawal threshold (g) response was then calculated from the resulting scores. Mechanical paw-withdrawal threshold was measured at baseline and at different time following treatments.

#### Cold hypersensitivity

Cold sensitivity was assessed using the acetone evaporation test. A single drop of acetone was applied to the plantar surface of the hind paw, and cold-evoked responses were quantified by measuring the total duration (seconds) of paw licking and lifting behaviors during a 5 min observation period. Cold sensitivity was measured at baseline and at different time following treatments.

#### Heat hyperalgesia

Mice were placed on a hot plate (Ugo Basile) set at 50 ± 0.1 °C. The latency to the first hind paw licking/withdrawal was taken as an index of the nociceptive threshold and detected before (basal) and after treatments. Cutoff time was set at 30 s. Heat hyperalgesia was measured at baseline and at different time following treatments.

### H_2_O_2_ assay

H_2_O_2_ levels were assessed in sciatic nerve tissue homogenates using the Amplex Red® assay (Invitrogen, Waltham, MA, USA), according to the manufacturer’s protocol. Fluorescence excitation and emission were measured at 540 and 590 nM, respectively. H_2_O_2_ production was calculated using the H_2_O_2_ standard and expressed as nmol/mg protein.

### Cell lines

Commercial human primary Schwann cells (hSCs, #P10351, Innoprot, Spain) were cultured and maintained in Schwann cell medium (#P60123, Innoprot, Spain) at 37 °C in 5% CO_2_ and 95% O_2_. Cells were routinely maintained until approximately 90% confluence and used for functional experiments. After 12 passages, cells were discarded and replaced. AAVpro-HEK293T cells (#632273, Takara, Diatech), were maintained in DMEM high glucose supplemented with heat inactivated FBS (10%), L-glutamine (4 mM),1 mM penicillin/streptomycin (1 mM) and sodium pyruvate (1 mM) at 37 °C in 5% CO_2_ and 95% O_2_.

### Plasmid constructs

pAAV[FLEXon]-CMV-rev(EGFP-5’-miR-30E-BfuAI-ORF-BfuAI-3’-miR-30E)-WPRE, obtained from Vector Builder, was used as backbone for a short hairpin RNA targeting gene of interest or Scrambled shRNA as negative control. shRNAs targeting the gene of interest (P1/P2 mNgfr, P3/P4 mNtrk1) were cloned by replacing the ORF with preannealed and phosphorylated oligonucleotides by using BfuAI compatible bases. All constructs were confirmed by Sanger sequencing. Primer sequences are listed in Supplementary Table 1.

### AAV production cell lysis and Iodixanol-based purification

AAVPro-HEK293T cells were plated in a CellBIND Polystyrene CellSTACK 2 Chamber (#3310, Corning, Corning, NY, USA) for 48 h to reach 80% confluence. Cells were washed with PBS, detached with trypsin-EDTA 0.05% (EuroClone, Milan, Italy). Cells (250 x 10^6^) were seeded in a CellBIND Polystyrene CellSTACK 5 Chamber (#3311; Corning) for 24 h, until 80% of confluence. Cells were transfected with 2.5 mg of DNA containing the three plasmids (packaging, helper and gene of interest (GOI)) in a 1:1:1 molar ratio. To produce rAAV that infects with high efficiency Schwann cells, Rep/Cap 2/rh10 was used (pAAV2/rh10 #112866, Addgene). To mainly infect primary sensor neurons, AAV2 rep-AAV-PHP.S were used (pUCmini-iCAP-PHP.S #103006, Addgene). Total DNA was diluted in 176 ml of OptiMEM (Thermo Fisher Scientific) and combined with PEI (1:3 DNA to PEI ratio) and mix. After 15 min of incubation, 350 mL of DMEM supplemented with FBS (2%) was added to the OptiMEM/DNA/PEI mix and used to replace the complete cells medium. Cells were maintained at 37 °C in 5% CO_2_ and 95% O_2_ for 72 h, before AAV particles were started to be collected. Cells were harvested and transferred into 50 mL conical tubes and centrifuged at 1000 × g for 10 min at 4°C. The supernatant was filtered through 0.45-μm PES membranes and then 25 mL of PEG solution (400 g of 40% polyethylene glycol + 24 g of NaCl in ddH2O to a final volume of 1.000 mL, pH 7.4) was added to every 100 mL of collected supernatant. The total solution was slowly stirred at 4 °C for 1 h and then, kept for 3 h without stirring at 4°C to allow full precipitation of particles. The solution was then centrifuged at 2.800 × g for 15 min at 4°C, the supernatant was discarded, and the virus was resuspended in a 10 ml of PBS/pluronic F68 (0.001%)/NaCl (200 mM) solution. Virus producing cell pellet was directly resuspended in 10 mL of PBS/Pluronic F68 (0.001%)/NaCl (200 mM) solution and cells were lysed by 4 cycles of freezing/thaw. Each cycle included a 30min step at −80°C, followed by a 10 min thaw at 37°C, interspersed with vortexing. Sample was then centrifuged at 3.200 × g for 15 min at 4°C and supernatant, containing the AAV particles was collected, while cell debris were discarded. Samples obtained by medium treatment and cell lyses, containing the AAV particles, were finally mixed, incubated with Benzonase (50 U/mL) at 37°C for 45 min to digest residual plasmids and residual genomic DNA / cellular RNA. Then, sample was centrifuged at 2.400 × g for 10 min at 4°C. The clarified supernatant was transferred to new tubes and was kept overnight at 4°C before purification.

Purification was carried out using a iodixanol gradient ultracentrifugation. Starting with a 60% iodixanol solution (OptiPrep; STEMCELL Technologies, Vancouver, Canada), a iodixanol gradient was prepared with a 15% solution [4.5 mL of iodixanol (60%) + 13.5 mL of NaCl/PBS-MK buffer (1M)], a 25% solution [5 mL of iodixanol (60%) + 7 mL of PBS-MK buffer (1×) + 30 μL of phenol red], a 40% solution [6.7 mL of iodixanol (60%) + 3.3 mL of PBS-MK buffer (1×)], and a 60% solution [10 mL of iodixanol (60%) + 45 μL of phenol red]. Each solution was added into a 39-mL Quick-Seal tube (Beckman Coulter, Brea, CA, USA) using 18 G needle syringe in the following order: 8 mL of the 15% iodixanol solution, 6 mL of the 25% iodixanol solution, 5 mL of the 40% iodixanol solution, and 5 mL of the 60% iodixanol solution. Finally, tubes were filled with the sample, sealed and, centrifuged in a Type 70 Ti rotor (Beckman Coulter) at 350.000 x g at 10°C for 90 min and then pierced with a 16 G needle on top and an 18 G needle at the interface between the 60% and 40% iodixanol gradients. Viral particles contained in the 40% iodixanol layer were fractioned in 1.5 mL microcentrifuge tubes and concentrated using Amicon Ultra-15 centrifugal filter units (molecular weight cut-off, 100 kDa; Merck Millipore). Before the concentration step, membranes were activated with 15 mL of 0.1% Pluronic F68 in PBS solution that were discarded and replaced with 15 mL of 0.01% Pluronic F68 in PBS solution. The tubes were centrifuged at 3.000 rpm for 5 min at 4 °C. The supernatant was discarded, and 15 mL of 0.001% Pluronic F68 in PBS + NaCl 200mM solution was added and centrifuged at 3.000 rpm for 5 min at 4 °C. The sample was then added and centrifuged at 3.500 rpm for 8 min at 4°C and flowthrough was discarded. During concentration process, formulation buffer (0.001% Pluronic F68 in PBS) was also added to the sample after few centrifugation steps to replace iodixanol and to avoid toxicity in animals after AAV injection. The viral title was quantified using RT-qPCR.

### Calcium imaging

Commercial primary hSCs were plated on poly-L-lysine-coated (8.3 µM) 35-mm glass coverslips and maintained at 37°C in 5% CO₂ and 95% O₂. Schwann cells were cultured for 24 h before calcium imaging experiments. On the day of experiments, cells were loaded (40 min) with Fura-2 AM-ester (5 µM) added to the buffer solution (37°C) containing (in mM): 2 CaCl₂, 5.4 KCl, 0.4 MgSO₄, 135 NaCl, 10 D-glucose, 10 HEPES and bovine serum albumin (BSA, 0.1%) at pH 7.4. Cells were then washed and transferred to a chamber mounted on the stage of a fluorescence microscope for recording (Axio Observer 7; equipped with a fast filter wheel, Digi-4 lens for excitation recording and Ultra- fast Sutter Lambda DG4 Xenon excitation source, range 300–700 nm; ZEISS, Stuttgart, Germany). For concentration-response experiments, commercial primary hSCs were exposed to recombinant NGF or proNGF (10 nM-100 µM), and intracellular calcium responses were recorded in real time for 10-20 min according to the experimental conditions. For pharmacological experiments, cells were stimulated with NGF (4 µM) or proNGF (2 µM) in the presence of thapsigargin (2 µM), Ca²⁺-free medium, LM11A-31 (1 µM), GW441756 (1 µM) or A967079 (30 µM). Control cells were treated with vehicle (0.001% DMSO). Results were expressed as the percent increase in fluorescence ratio 340/380 (R_340/380_) over baseline normalized to the maximum effect induced by ionomycin (5 µM) added at the end of each experiment. For selected experiments, calcium responses were quantified as area under the curve (AUC).

### Mitochondrial ROS imaging

Mitochondrial ROS production was evaluated using MitoSOX Red mitochondrial superoxide indicator (#M36008; Thermo Fisher Scientific, Waltham, MA, USA). Commercial primary hSCs were plated on poly-L-lysine-coated (8.3 µM) 35-mm glass coverslips and maintained at 37°C in 5% CO₂ and 95% O₂. Cells were cultured for 24 h before experiments. On the day of imaging, cells were incubated with MitoSOX Red (5 µM) for 10 min at 37°C in the dark and subsequently washed before fluorescence acquisition. Cells were transferred to the imaging chamber mounted on the fluorescence microscope (Axio Observer 7; ZEISS, Stuttgart, Germany). Commercial primary hSCs were exposed to NGF (4 µM) or proNGF (2 µM) in the presence of LM11A-31 (1 µM), GW441756 (1 µM), MitoTEMPO (5 µM) or vehicle (0.001% DMSO). MitoSOX Red fluorescence changes were monitored in real time for approximately 15 minutes. Results were expressed as relative fluorescence changes over baseline and quantified as area under the curve (AUC) of the fluorescence response.

### H₂O₂ imaging

A genetically encoded probe for intracellular hydrogen peroxide (HyPer7.2; (Pak *et al*, 2020)) was used for real-time imaging of H₂O₂ production in live commercial primary hSCs. Cells were plated on poly-L-lysine-coated (8.3 µM) 35-mm glass coverslips and transfected with HyPer7.2 DNA (2 µg) using jetOPTIMUS® DNA transfection reagent (#55-250; Polyplus, Lexington, MA, USA) according to the manufacturer’s instructions. After 24–48 h, cells were washed and transferred to a chamber mounted on the stage of a fluorescence microscope for recording (Axio Observer 7; equipped with fast filter wheel and Digi-4 lens for excitation recording; ZEISS, Stuttgart, Germany). Commercial primary hSCs expressing HyPer7 were exposed to NGF (4 µM) or proNGF (2 µM), or vehicle (0.001% DMSO), and intracellular H₂O₂ variations were monitored for approximately 10 minutes. To investigate the signaling pathways involved in NGF- and proNGF-induced oxidative responses, experiments were performed in the presence of LM11A-31 (1 µM), ML-171 (1 µM), MitoTEMPO (5 µM), GW441756 (1 µM) or A967079 (30 µM). HyPer7 fluorescence changes were expressed as variations over baseline and quantified as area under the curve (AUC) of the fluorescence response over time.

### Data and statistical analysis

The results are expressed as the mean ± SEM. For multiple comparisons, a one-way ANOVA followed by a post-hoc Bonferroni’s test was used. The two groups were compared using Student’s t-test. For behavioral experiments with repeated measures, a two-way mixed-model ANOVA followed by a post-hoc Bonferroni’s test was used. Statistical analyses were performed on raw data using GraphPad Prism 8 (GraphPad Software Inc.). EC_50_ and EC_80_, values were determined from non-linear regression models using Graph Pad Prism 8. P-values less than 0.05 (P < 0.05) were considered significant.

## Data availability

All data generated for this study are available upon request.

